# A custom genotyping array reveals population-level heterogeneity for the genetic risks of prostate cancer and other cancers in Africa

**DOI:** 10.1101/702910

**Authors:** Maxine Harlemon, Olabode Ajayi, Paidamoyo Kachambwa, Michelle S. Kim, Corinne N. Simonti, Melanie H. Quiver, Desiree C. Petersen, Anuradha Mittal, Pedro Fernandez, Ann W. Hsing, Shakuntala Baichoo, Ilir Agalliu, Mohamed Jalloh, Serigne M. Gueye, Nana Yaa Snyper, Ben Adusei, James E. Mensah, Afua O.D. Abrahams, Akindele O. Adebiyi, Akin Orunmuyi, Oseremen I. Aisuodionoe-Shadrach, Maxwell M. Nwegbu, Maureen Joffe, Wenlong C. Chen, Hayley Irusen, Alfred I. Neugut, Yuri Quintana, Moleboheng Seutloali, Mayowa Fadipe, Christopher Warren, Marcos H. Woehrmann, Peng Zhang, Chrissie Ongaco, Michelle Mawhinney, Jo McBride, Caroline Andrews, Marcia Adams, Elizabeth Pugh, Timothy R. Rebbeck, Lindsay Petersen, Joseph Lachance

**Author notes:** **Corresponding author**, Joseph Lachance, 950 Atlantic Dr. Atlanta, GA 30332.

## Abstract

Although prostate cancer is the leading cause of cancer mortality for African men, the vast majority of known disease associations have been detected in European study cohorts. Furthermore, most genome-wide association studies have used genotyping arrays that are hindered by SNP ascertainment bias. To overcome these disparities in genomic medicine, the <u>M</u>en of <u>A</u>frican <u>D</u>escent and <u>Ca</u>rcinoma of the <u>P</u>rostate (MADCaP) Network has developed a genotyping array that is optimized for African populations. The MADCaP Array contains more than 1.5 million markers and an imputation backbone that successfully tags over 94% of common genetic variants in African populations. This array also has a high density of markers in genomic regions associated with cancer susceptibility, including 8q24. We assessed the effectiveness of the MADCaP Array by genotyping 399 prostate cancer cases and 403 controls from seven urban study sites in sub-Saharan Africa. We find that samples from Ghana and Nigeria cluster together, while samples from Senegal and South Africa yield distinct ancestry clusters. Using the MADCaP array, we identified cancer-associated loci that have large allele frequency differences across African populations. Polygenic risk scores were also generated for each genome in the MADCaP pilot dataset, and we found that predicted risks of CaP are lower in Senegal and higher in Nigeria.

**Significance:** We have developed an Africa-specific genotyping array which enables investigators to identify novel disease associations and to fine-map genetic loci that are associated with prostate and other cancers.

## Introduction

Prostate cancer (CaP) is a complex disease that disproportionally affects men of African descent (1). CaP is the leading cause of cancer-related mortality in African men (2). In the United States, African American men have a higher risk of developing CaP, and an even higher increased risk of dying from it compared to men of European or Asian descent (3). In the United Kingdom, men of African descent have an increased risk of being diagnosed and dying from CaP (4). Furthermore, the highest reported mortality rates of CaP are found in the Caribbean men of African descent (5). Multiple socioeconomic, environmental, and genetic factors contribute to this health inequity.

Cancer is considered a genetic disease, and CaP has a high heritability: on the order of 58% (6). Risks of CaP tend to run in families; the relative risk of men with affected fathers is 2.1-fold higher compared to men without a family history and the relative risk of men with affected brothers is 2.3-fold higher compared to men without family history (7). In recent years, multiple genome-wide association studies (GWAS) have detected genetic associations with CaP (8–11). Collectively, these studies have yielded over 200 independent CaP risk-associated loci, and one key genomic region that has been repeatedly implicated in CaP and other cancers is 8q24 (12–14). The most comprehensive CaP GWAS analyzed to date included more than 140,000 cases and controls of European ancestry (10). In this study, Schumacher et al. generated a polygenic risk score (PRS) that successfully classified individuals into high and low risk categories.

Unfortunately, most GWAS have not focused on populations from sub-Saharan Africa. As of 2016, 81% of all GWAS samples were of European ancestry and 14% of all GWAS samples were of East Asian ancestry (15). Additionally, existing genotyping arrays do not adequately capture genetic variation in diverse African populations. Both of the aforementioned issues limit what is known about cancer genetics in African populations. Thus, the current set of known disease-associated loci is enriched for alleles with an intermediate frequency in Europe or Asia, but not Africa. This lack of representation can exacerbate existing health disparities by not capturing relevant genetic risk associations in African populations (16,17). Disease associations do not always replicate across populations and ancestries, and the directions of risk associations at cancer-associated loci may differ across study populations (18,19). Previous work also indicates that risks of CaP vary by genetic ancestry across the globe (20,21). For example, an earlier GWAS in Ghana showed that the most promising SNP in this African population has not been identified in other populations (22). Hence, there is a clear need for more studies that analyze the genetics of African populations (23,24). Commonly used genotyping arrays tend to use markers that were originally ascertained in European populations (25). This can cause polygenic risks of complex diseases, including CaP, to be wrongly estimated (26). For example, the OncoArray Consortium has developed an array with over 500,000 markers, half of which are in genomic regions that tag cancer susceptibility (28). However, the OncoArray is not enriched for African polymorphisms. By contrast, the H3Africa Consortium has developed an array that includes over two million markers (27), but the H3Africa Array was not specifically designed for cancer studies. Existing arrays may therefore be sub-optimal for detecting cancer associations in African populations.

To address this problem, the <u>M</u>en of <u>A</u>frican <u>D</u>escent and <u>Ca</u>rcinoma of the <u>P</u>rostate (MADCaP) Network (29) developed a customized genotyping array optimized for fine-mapping and detecting novel associations with CaP in African populations. The MADCaP array will ultimately be used in an African GWAS containing over 6000 cases and controls. Here, we analyze a pilot dataset of over 800 individuals from sub-Saharan Africa. In this paper, we compare multiple genotyping platforms and test the efficacy of the MADCaP Array by genotyping samples from seven African study sites. Using data derived from the MADCaP Array, we also infer population structure, identify cancer-associated loci that have large allele frequency differences across Africa, and quantify polygenic risks of CaP in urban African populations.

## Materials and Methods

### Inclusion criteria for markers on the MADCaP Array

The MADCaP Array was developed using the Applied Biosystems™ Axiom™ genotyping solution from Affymetrix/ThermoFisher Scientific. This array consists of a two-peg design. Multiple inclusion criteria were used for markers on the MADCaP Array, including: enrichment for GWAS loci, markers near cancer susceptibility loci, prostate eQTLs (expression quantitative trait loci), markers found on other arrays, and markers tagging African polymorphisms. 38,649 unique markers that are associated with traits and diseases from the NHGRI-EBI GWAS Catalog are included on the MADCaP Array (30). Using 1000 Genomes Project (31) data, we included every SNP with an African minor allele frequency (MAF) > 0.05 that was located within 50kb of a known CaP hit or within 5kb of other cancer associations. We used the Genotype-Tissue Expression (GTEx V7) project (32) to identify SNPs that modify gene expression in the prostate (i.e. prostate eQTLs). The MADCaP Array contains a total of 24,595 prostate eQTLs (p-value cutoff for inclusion: 10^-9^). Markers were also preferentially included if they overlapped the OncoArray or H3Africa Array. Working with ThermoFisher Scientific, a GWAS backbone was built using Applied Biosystems Axiom genotyping array technology by iteratively selecting markers that maximized the ability to impute African genetic variation. When possible, we used probes that had a prior track record of working on existing genotyping arrays. Multiple probes per marker were included for CaP loci and unvalidated markers. An overlap of more than 1000 markers were chosen to be on both pegs, with priority given to CaP loci and markers satisfying multiple inclusion criteria. Table S1 lists successfully called markers on the MADCaP Array as well as inclusion criteria details.

### Assessment of Imputation performance

Imputation performance of the MADCaP Array was computed using the African Genome Resource reference panel, comprising of data from 4,956 individuals of African descent (33). African polymorphisms with MAF > 0.05 were classified as common SNPs and African polymorphisms with a MAF between 0.01 and 0.05 were classified as rare SNPs. Imputation was performed with IMPUTE2 (v2.3.2) software using 10-fold cross validation (34). Coverage in each population was calculated as the proportion of polymorphisms in the African Genome Resource reference panel in high LD (r^2^ ≥ 0.8) with markers on the MADCaP Array.

### Biospecimen and DNA quantification

Biospecimens were obtained with informed consent using protocols approved from each study site’s Institutional Review Board/Ethics Review Board. Blood samples were collected in EDTA vacutainer tubes and stored at either −20°C or −80°C dependent on the timeframe for DNA extraction. DNA was isolated using QIAamp DNA Blood kits. A total of 1.8 to 3.0 μg high purity DNA at a concentration of 30 to 50 ng/μl per sample was submitted for genotyping. DNA was transferred from study sites to genotyping laboratories using BioMatrica DNAStable 2D barcoded plates. Samples were then rearrayed into plates using a BioMicroLab XL20 at a minimum concentration of 10 ng/μl in 50μl. All samples were run on the Infinium QC array and the MADCaP Array. Plate maps used a randomized block design to control for study site and case vs. control status.

### SNP calling, QC, and data curation

Standard quality control (QC) procedures for Axiom genotyping data analysis were performed (35,36). Sample pre-processing was performed according to guidelines provided in the Thermo Fisher Scientific Axiom Genotyping Solution Data Analysis Guide (36). The custom MADCaP Array is based on a two-peg design. Peg 1 contains a total of 852,610 probe sets, covering 801,275 markers. Peg 2 contains a total of 790,524 probe sets, covering 790,170 markers. 1,902 probe sets overlap both pegs. Raw data CEL files, representing more than 802 samples, as well as 28 technical replicates and additional controls, were imported into the Axiom Analysis Suite (AxAS) version 4.0.3.3 for filtering of sample call rate and clustering of SNP genotype calls. Samples with DishQC ≥ 0.82 and a QC call rate >97% were included for downstream genotyping analysis. Additional SNP metrics included tests of Hardy-Weinberg proportions, reproducibility of genotyping calls, and identifying Mendelian inconsistencies. The Centre for Proteomics and Genomics Research (CPGR) in Cape Town, South Africa and the Center for Inherited Disease Research (CIDR) at Johns Hopkins University independently assessed QC metrics for each probe set.

Data from both pegs of the MADCaP Array were merged and PLINK was used to remove related samples (identity-by-state > 0.5). Multi-allelic SNPs were excluded from downstream analyses. After filtering markers with low call rates and excluding poorly called and related samples, 1,513,172 markers and 802 samples were used in subsequent population genetic analyses. This MADCaP pilot dataset contains 399 CaP cases and 403 controls. Details of MADCaP case and control recruitment have been previously reported (29).

### Array comparisons

We compared markers on arrays developed by the MADCaP Network (29), the OncoArray Consortium (28), and the H3Africa Consortium (27). Genomic positions from the MADCaP Array, Infinium Oncoarray and H3Africa Array were intersected to determine overlapping markers between arrays. The *LiftOver* bioinformatics tool was used to ensure that all genomic positions used build GRCh38/hg38 of the human reference genome. Mean derived allele frequencies (DAF) for each array were calculated as described previously (26). This involved obtaining allele frequencies from each of the five continental regions using the 1000 Genomes Project (31): Africa (AFR), Americas (AMR), East Asia (EAS), Europe (EUR), and South Asia (SAS). Calculations of mean derived allele frequencies used 450,000 markers that were randomly selected, without replacement, from each array. The joint allele frequency distribution of all 1,513,172 markers on the MADCaP Array was found by comparing African and pooled non-African data from the 1000 Genomes Project.

### MDS and ADMIXTURE

Using PLINK, an LD pruned subset of 25,000 autosomal SNPs for each of 802 samples was obtained (MAF > 0.05, r^2^ < 0.8). The same subset of SNPs was used for Multidimensional Scaling (MDS) and ADMIXTURE analyses. Two dimensional MDS plots were generated using PLINK and R. ADMIXTURE software (37) was run for K = 2 through K = 5. Cross-validation was performed to determine the optimal K value.

### Runs of homozygosity and LD decay

As per Schlebusch et al. (38), runs of homozygosity were identified using PLINK for homozygous lengths between 500kb and 1000kb. This analysis was repeated for all 802 samples in the MADCaP dataset. Individual runs of homozygosity were summed to yield the cumulative runs of homozygosity (cROH) for each sample. For each MADCaP study site, PLINK v1.90b6.9 was used to calculate LD between all variants with a MAF > 0.10. These calculations were made for all pairs of markers within 100kb and 100 marker windows. Distances between genetic variants were used to place pairs of variants into 1 kb bins. For each study site, the mean r^2^ between pairs of genetic variants was calculated for each 1 kb bin.

### Identification of divergent loci via PBS calculations

Population branch statistics (PBS) were calculated as per Yi et al. (39). Data from multiple MADCaP study sites were pooled to yield allele frequencies for three populations: Senegal (HOGGY), Ghana & Nigeria (37 Military, KBTH, UATH, and UCH) and South Africa (WITS and SU). Genetic distances between pairs of populations were calculated using Weir and Cockerham’s F_st_ (40). The following equations were then used to calculate PBS scores for three different evolutionary branches:

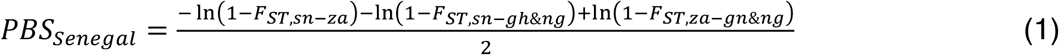

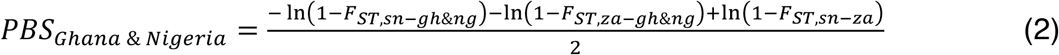

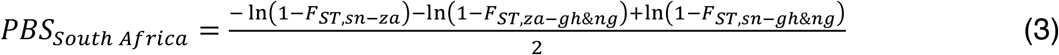

Subscripts in the above equations refer to country codes: *sn* for Senegal, *gh* for Ghana, *ng* for Nigeria, and *za* for South Africa. Undefined and negative values of Weir and Cockerham’s F_st_ were treated as zero in the above equations, and undefined or negative PBS scores were also treated as zero. PBS scores were calculated for 2,477 unique markers from the NHGRI-EBI GWAS Catalog (30) that yield 5,337 cancer and cancer-related associations.

### Calculation of polygenic risk scores (PRS)

Polygenic risk scores (PRS) were built using a curated set of 141 CaP-associated loci. Schumacher et al. previously developed a 147 loci PRS for CaP (10), and 119 of these 147 markers are on the MADCaP Array. Proxies were found for 22 of the remaining 27 markers, by identifying markers on the MADCaP array in LD with loci from the Schumacher PRS (r^2^ > 0.4). LDlink (41) was then used to select alleles that tag increased CaP risk at proxy markers. Table S2 lists markers that were used to generate the PRS described here.

As per Schumacher et al. (10), effect size information was incorporated into PRS calculations. For each locus, we counted whether an individual has 0, 1, or 2 copies of the risk-increasing allele (i.e. the allele dose *g_ij_* for locus *i* in individual *j*). Here we used adjusted effect sizes: 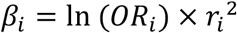, where effect sizes from Schumacher et al. (10) are scaled by how well proxy markers tag each disease-associated locus. Doses of risk-increasing alleles were weighted by adjusted effect sizes and summed across all 141 loci to a raw PRS for each individual.

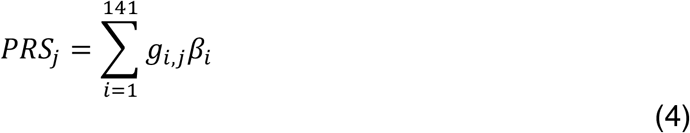

PRS were calculated for 802 MADCaP samples and 240 men of European ancestry from 1000 Genomes Project (31). Standardized PRS values were then generated for all 1,042 individuals by scaling raw PRS values to have a mean of zero and standard deviation of one.

## Results

### Imputation using the MADCaP Array

Using whole genome sequences, we quantified the extent to which the MADCaP array tags African genetic variation (Fig. 1). Depending on the population, 94% to 99% of common African SNPs were successfully tagged by the MADCaP Array (r^2^ ≥ 0.8, MAF > 0.05). The MADCaP Array also tagged 63% to 97% of rare African SNPs (r^2^ ≥ 0. 8, MAF between 0.01 and 0.05). It captures a larger fraction of Ugandan genetic variation than Ethiopian and KhoeSan variation. Fig. 1 also shows that the MADCaP Array successfully tags variation in admixed African-Caribbean and African-American genomes. Regardless of population, the MADCaP Array successfully captures a large fraction of African genetic variation.

**Figure 1.**
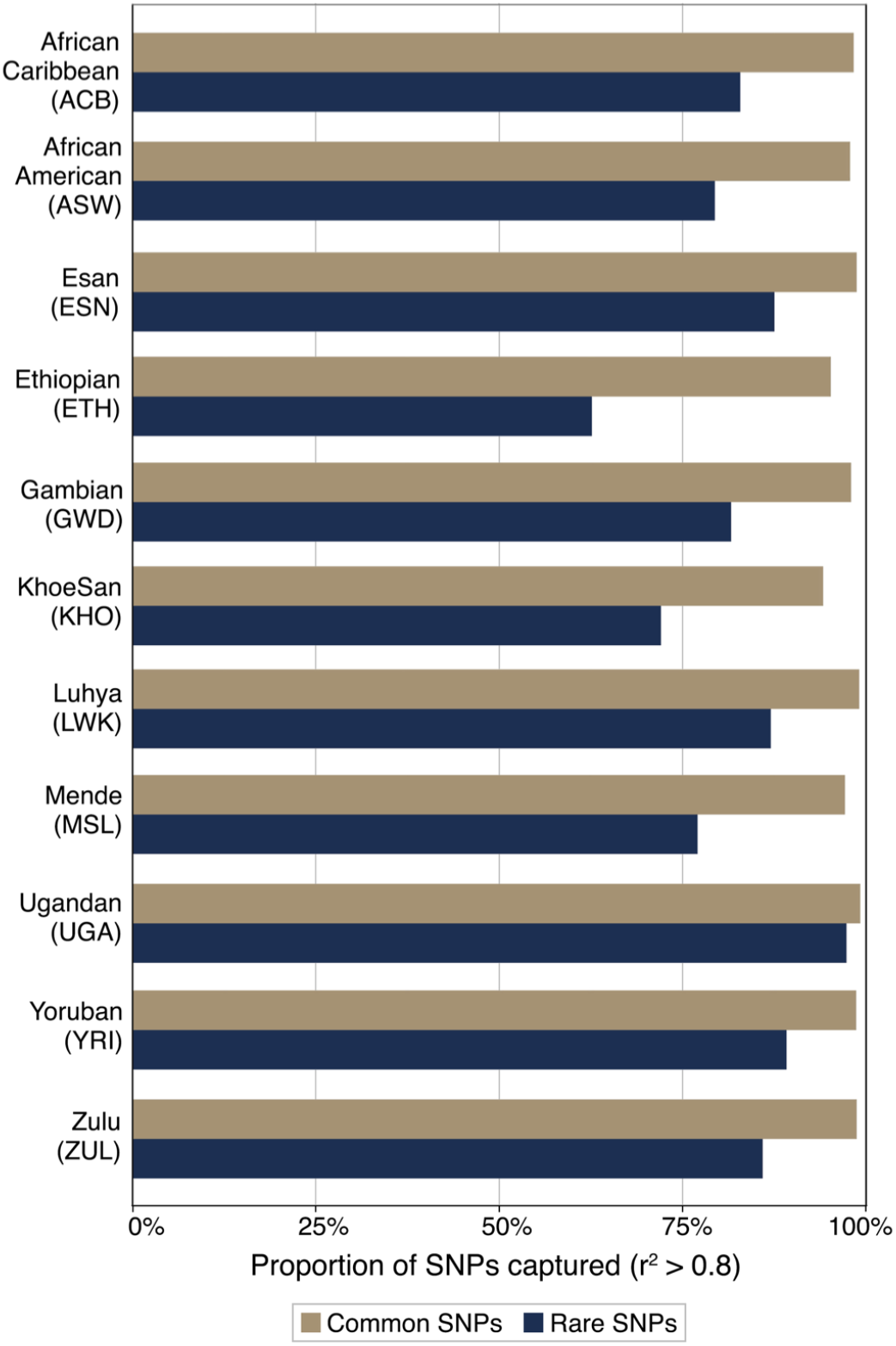
African SNPs are accurately imputed using MADCaP Array. Proportions of SNPs in 11 populations of African ancestry that are successfully tagged by markers on the MADCaP Array are shown (r^2^ ≥ 0.8). Here, common SNPs have a MAF > 0.05 and rare SNPs have a MAF between 0.01 and 0.05.

### Comparisons with other arrays

Many of the markers on the MADCaP Array are shared with the Infinium OncoArray and the H3Africa Array (Fig. 2A). Overall, 73,019 markers are included on all three arrays. A total of 131,469 markers are shared between the MADCaP Array and the OncoArray, and a total of 398,460 markers are shared between the MADCaP Array and the H3Africa Array. This overlap will facilitate data harmonization and the ability to combine genotype information from different arrays into the same study.

**Figure 2.**
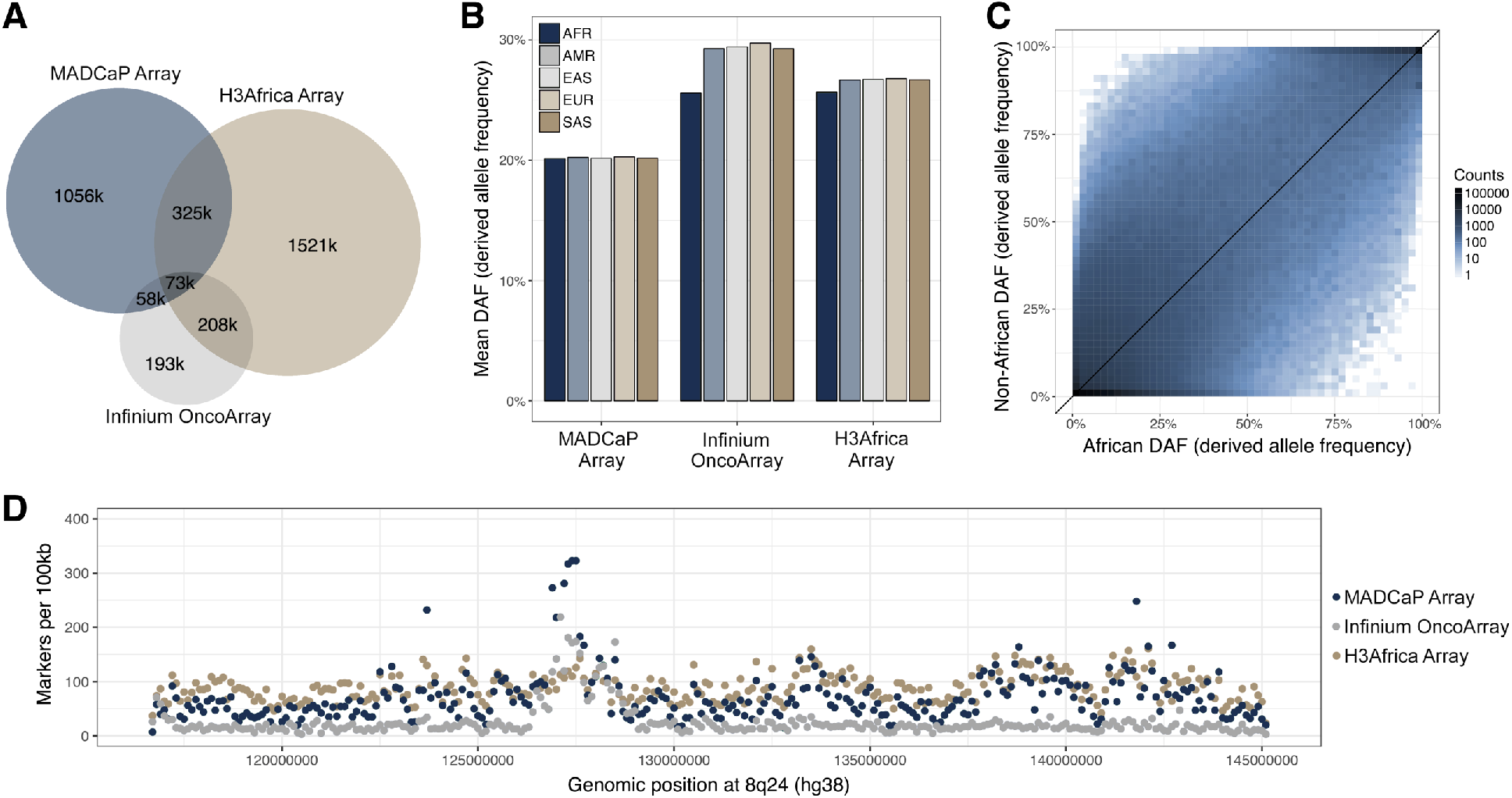
Comparisons between the MADCaP Array, Infinium OncoArray, and H3Africa Array. **A**, Venn Diagram showing overlap between markers on each array. Sizes of circles are proportional to the number of markers on each array. **B**, Mean DAF of markers on the MADCaP Array, Infinium OncoArray, and H3Africa Array. Continental allele frequencies from the 1000 Genomes Project are shown here. **C**, Joint site frequency spectrum of markers on the MADCaP Array. African and pooled non-African allele frequencies from 1000 Genomes Project are shown here. Shading indicates the number of markers on the MADCaP array that are in each bin. **D**, Density of markers per non-overlapping 100kb window. The 8q24 genomic region is shown here.

We compared the mean derived allele frequencies (DAF) of markers found on the MADCaP, OncoArray, and H3Africa arrays, using Continental allele frequencies from the 1000 Genomes Project (Fig. 2B). The null expectation here is that the mean DAF should be the same for each population since all humans are evolutionarily equidistant to other primates. Mean DAFs of markers on the MADCaP Array were similar for each continental population, suggesting that the MADCaP Array is relatively unbiased with respect to SNP selection. By contrast, the mean DAFs of markers on the OncoArray and H3Africa Array were lower for African populations than non-African populations (Fig. 2B). These DAF differences are indicative of SNP ascertainment bias (26). Examining the joint site frequency spectrum of non-African and African populations, similar counts of MADCaP markers are found above and below the diagonal in Fig. 2C. One exception to this pattern is that the MADCaP Array is enriched for markers that are polymorphic in Africa but monomorphic outside of Africa, but not vice-versa.

Densities of markers that are found in different genomic regions vary by genotyping array. Here, we focus on 8q24, a cancer-associated genomic region that contains *PCAT2, CCAT2,* and the proto-oncogene *c-myc.* Numbers of markers per 100kb are shown for three different arrays in Fig. 2D. The MADCaP Array contains a moderately high density of markers across the genome, with peaks near known cancer-associated loci. Neighboring markers on the MADCaP Array have a median distance of 856bp and a mean distance of 2082bp. The Infinium OncoArray has high marker densities near cancer-associated loci, but a low density of markers for other parts of the genome. In contrast, the H3Africa Array has a moderately even density of markers across the entire genome.

### Efficacy of the MADCaP Array

We tested the efficacy of the MADCaP Array by genotyping over 800 African individuals from seven MADCaP study sites (Fig. 3A): the Hôpital Général de Grand Yoff/Institut de Formation et de Recherche en Urologie in Dakar, Senegal (HOGGY), 37 Military Hospital in Accra, Ghana (37 Military), Korle-Bu Teaching Hospital in Accra, Ghana (KBTH), University College Hospital in Ibadan, Nigera (UCH), University of Abuja Teaching Hospital in Abuja, Nigeria (UATH), WITS Health Consortium/National Health Laboratory Services in Johannesburg, South Africa (WITS), and Stellenbosch University in Cape Town, South Africa (SU). Sample accrual was restricted to individuals with sub-Saharan African ancestry; admixed individuals with European ancestry from Cape Town were excluded. Of the MADCaP samples analyzed herein, 399 are CaP cases and 403 are controls (Fig. 3B).

**Figure 3.**
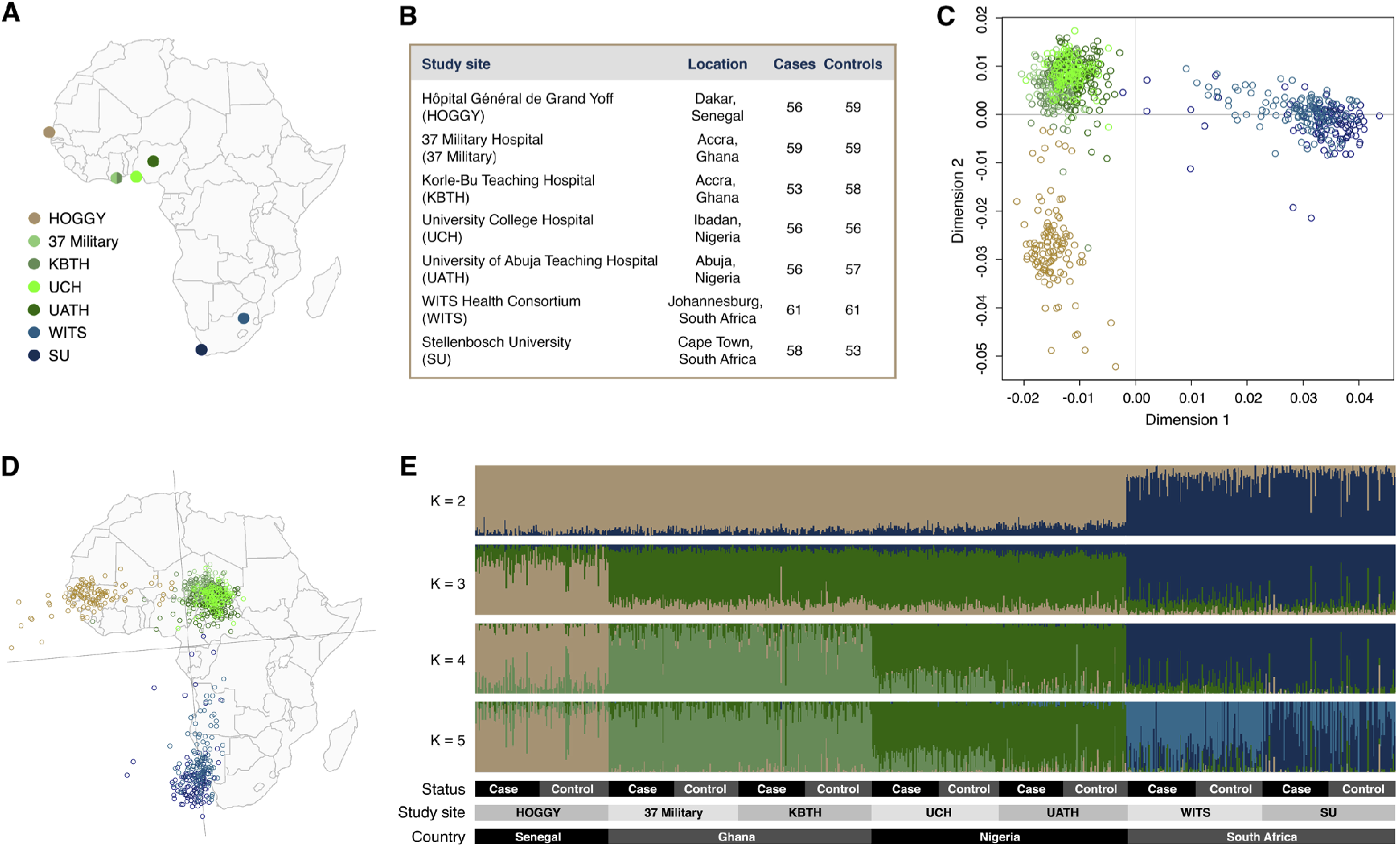
The MADCaP Array reveals population structure and shared genetic ancestries among urban African study sites. **A**, Geographic locations of each MADCaP study site. **B**, Numbers of cases and controls from each study site. **C**, Two-dimensional MDS plot of 802 MADCaP samples. Senegalese samples are represented by gold circles, Ghanaian and Nigerian samples are represented by green circles, and South African circles are represented by blue circles. D, Genes mirror geography when the two-dimensional MDS plot is rotated clockwise. **E**, ADMIXTURE plot of 802 MADCaP samples. The best to genetic data occurs at K = 3.

Up to 94.9% of the markers on peg 1 and 95.9% of the markers on peg 2 passed QC filtering. We note that probe sets from the MADCaP Array were 2.4 times less likely to fail than probe sets from the OncoArray Array (28). For both peg 1 and peg 2, mean call rates, reproducibility, and concordance all exceeded 99.5%, and only a small subset of markers had Mendelian inconsistencies (Table 1). Overall, we find that the MADCaP Array is an effective genotyping platform.

**Table 1.** Genotyping metrics for peg 1 and peg 2 of the MADCaP Array.

| QC metric | MADCaP Array<br>(Peg 1) | MADCaP Array<br>(Peg 2) |
| --- | --- | --- |
| Mean call rate<br>(proportion markers called) | 99.55% | 99.63% |
| Mean reproducibility<br>(Same calls for each probe) | 99.85% | 99.90% |
| Mean concordance<br>(Same calls in replicates) | 99.53% | 99.56% |
| Error rate<br>(Mendelian inconsistencies) | 0.051% | 0.032% |

### Population structure and genetic admixture

We used two-dimensional multidimensional scaling (MDS) plots to detect population structure among MADCaP samples and study sites. Individuals who have similar genomes are located close to one another in MDS space. MADCaP samples fall into three broad clusters in Fig. 3: Senegalese individuals (gold) are found in the bottom left, Ghanaian and Nigerian individuals (green) are found in the top left, and South African individuals (blue) are found in the top right. Nigerians from Ibadan (UCH, light green) are closer in MDS space to Ghanaian individuals than Nigerians from Abuja (UATH, dark green). The right-to-left gradient of blue points in MDS space suggest that some individuals from South Africa share a fraction of their genetic ancestry with present-day Nigerians. Rotating the MDS plot 85 degrees clockwise reveals that genes mirror geography, at least for the African populations analyzed in our study (Fig. 3D). Samples from geographically close locations tend to share greater amounts of genetic similarity.

ADMIXTURE plots reveal shared ancestry among MADCaP samples (Fig. 3E). In these plots, individuals are linear mixtures of multiple genetic ancestries – indicated by different colors. Cross-validation error is minimized at K= 3, i.e. the best fit to the data occurs for three ancestry colors (Fig. S1). At K = 2, we are able to distinguish between West African and South African populations. Setting K = 3 reveals three major ancestry clusters: gold in Senegal, green for Ghana and Nigeria, and blue in South Africa. At K = 4 ancestry patterns match each country. Intriguingly, individuals from Ibadan, Nigeria (UCH) share ancestry with samples from Ghana, i.e. they contain moderate amounts of light green ancestry at K = 4. Similarly, individuals from Johannesburg, South Africa contain traces of genetic ancestry that are primarily found in Nigeria (dark green), perhaps due to the Bantu expansion during the last 5,000 years (42). K = 5 reveals evidence of population structure within South Africa, with greater proportions of light blue ancestry found in Johannesburg (WITS) compared to Cape Town (SU). Both study sites from Accra, Ghana (37 Military and KBTH) have similar genetic ancestry profiles. Finally, we note that cases and controls for each study site are ancestry-matched. On a genome-wide scale, individuals in the MADCaP study with CaP have similar ancestry proportions compared to healthy MADCaP controls.

### Runs of homozygosity and linkage disequilibrium

Genotyping arrays can be used to identify runs of homozygosity – stretches of DNA where maternally and paternally inherited haplotypes are identical. Using the MADCaP array, we quantified cumulative runs of homozygosity (cROH) in each genome (Fig. 4A). Although there is heterogeneity within each study site, cROH are smaller for South African genomes than Senegalese, Ghanaian, or Nigerian genomes analyzed in this study (p-value = 3.31 × 10^-9^, Wilcoxon rank-sum tests). This lower homozygosity can either be due to large historical population sizes or due to admixture.

**Figure 4.**
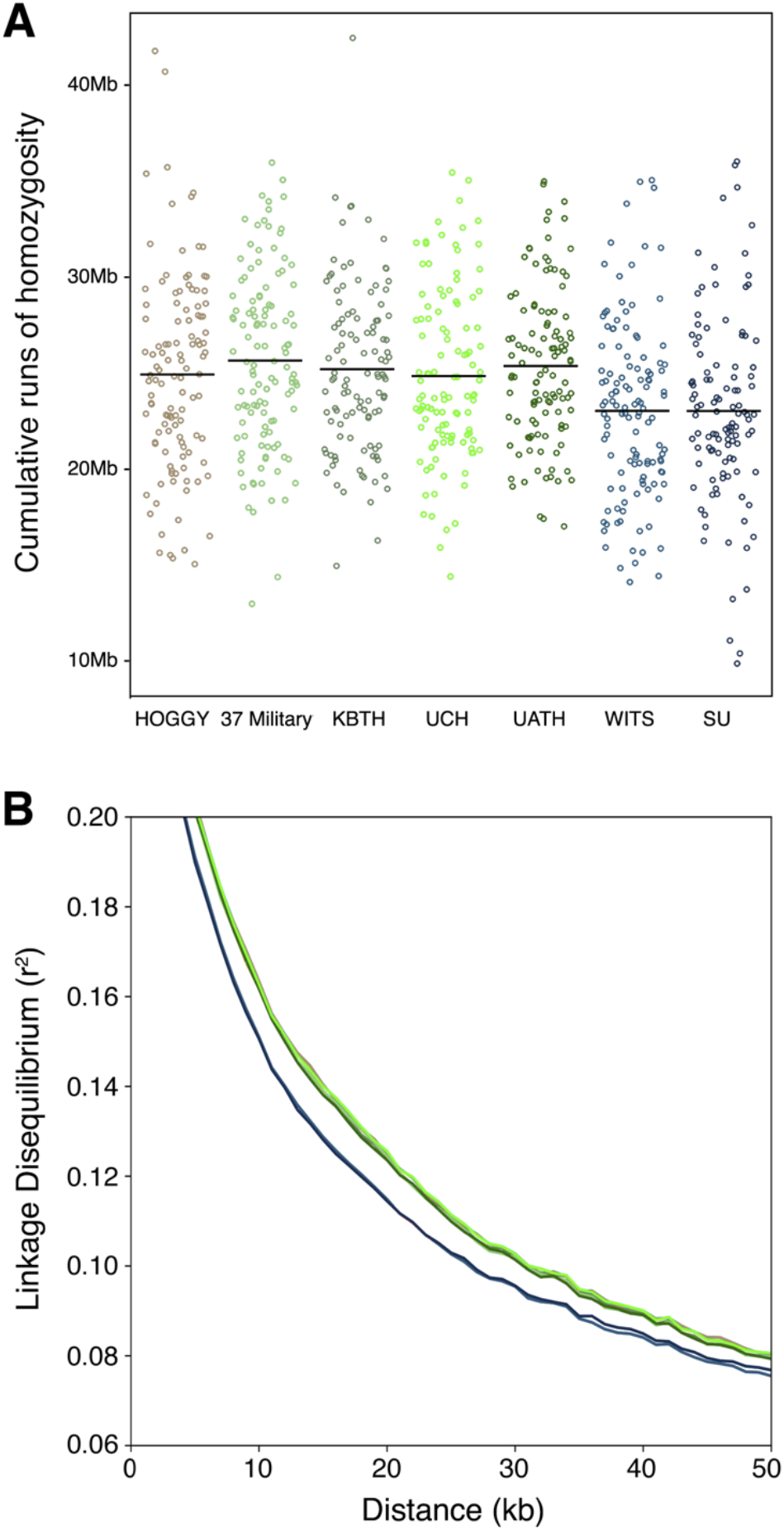
Runs of homozygosity and LD decay curves vary by African study site. **A**, Cumulative runs of homozygosity (cROH) 500kb to 1000kb in length for each MADCaP sample, labelled by study site. **B**, LD decay curves for each study site. Gold indicates Senegalese data, green indicates Ghanaian and Nigerian data, and blue indicates South African data. South African study sites have less LD than West African study sites (Senegalese data overlaps Ghanaian and Nigerian data).

To distinguish whether this lower homozygosity is due to large historical population sizes or admixture, we calculated LD decay curves for each of the seven MADCaP study sites (Fig. 4B). Populations with small effective population sizes have more LD than populations with large effective population sizes (43). Admixture also increases the amount of LD (44). In general, we observed less LD for South African sites than other study sites (WITS and SU in Fig. 4B). These differences in LD decay curves do not appear to be due to admixture, since the South African populations studied here have similar levels of admixture to other African populations (Fig. 3E). Overall, the data in Fig. 4 support the idea that historic population sizes were larger in South Africa than West Africa. One implication of the smaller haplotype blocks that are found in genomes from Johannesburg and Cape Town is that GWAS using these samples will require arrays with high densities of markers, a characteristic that is shared by the MADCaP Array.

### Divergent allele frequencies at cancer-associated loci

Risk allele frequencies at cancer-associated loci can vary across the African continent. To identify genetic variants that have large allele frequency differences across MADCaP populations, population branch statistic (PBS) scores were calculated for all observed cancer-associated loci in the NHGRI-EBI GWAS Catalog that have markers on the MADCaP Array, as well as loci associated with other cancer-related traits (e.g., skin pigmentation and smoking). These scores were calculated for three different evolutionary branches: Senegal (Fig. 5A), Ghana & Nigeria (Fig. 5B), and South Africa (Fig. 5C). Here, CaP hits used in PRS calculations are represented by black points, while gray and colored points indicate other cancer-associated loci. Table S2 contains PBS scores and allele frequencies for 141 CaP markers used in MADCaP PRS calculations. Table S3 contains PBS scores and allele frequencies for 2,477 markers that are associated with cancer and cancer-related traits.

**Figure 5.**
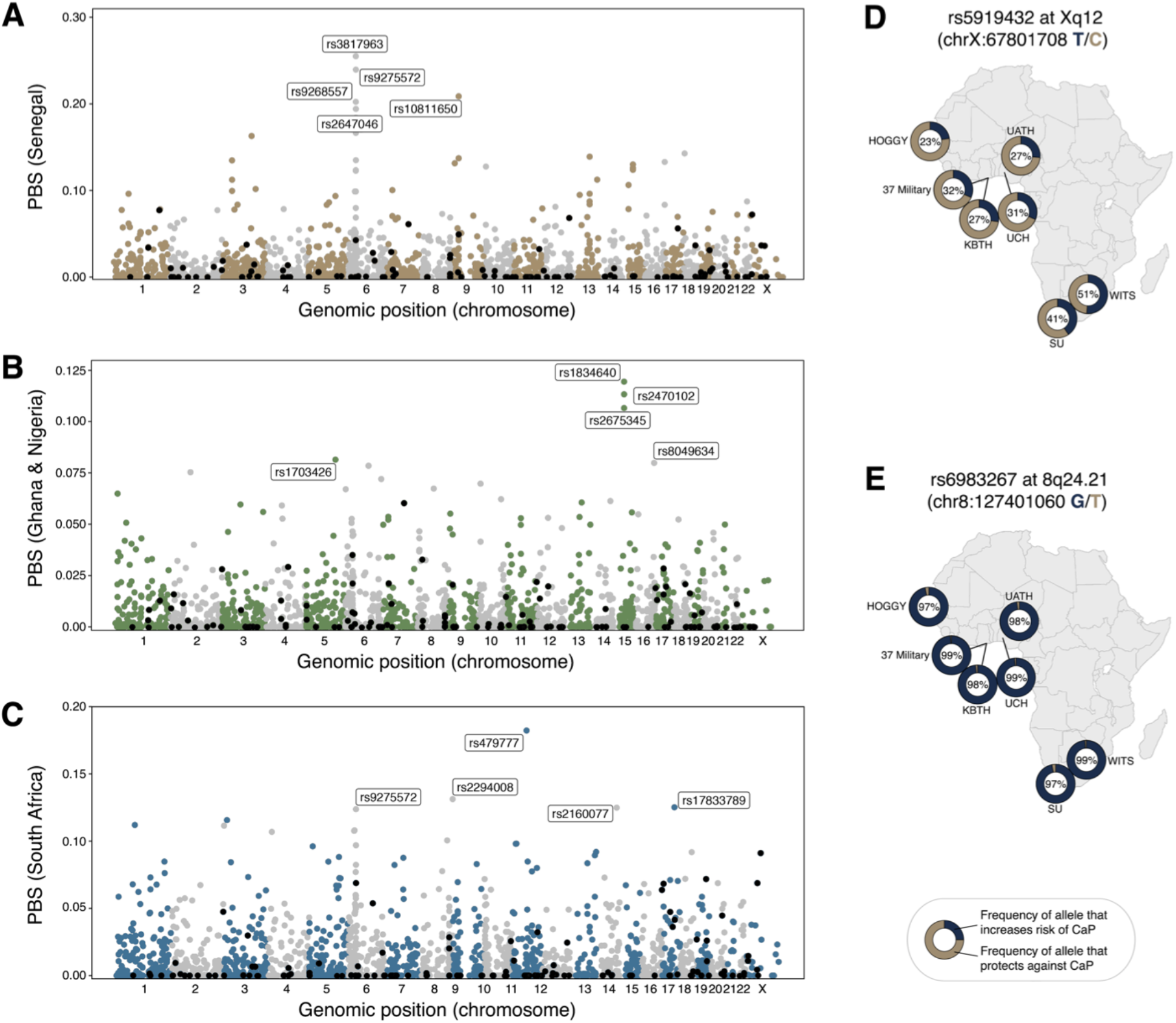
PBS scores identify divergent loci in Africa that are associated with cancer risks. CaP associations from the Schumacher et al. 2018 GWAS (10) are represented by black points, and other cancer-associated loci are represented by gray and colored points. **A**, PBS scores for the Senegal branch. **B**, PBS scores for the Ghana & Nigeria branch. **C**, PBS scores for the South African branch. **D**, Allele frequencies at the CaP-associated SNP rs5919432 vary greatly across Africa. **E**, Allele frequencies at the CaP-associated SNP rs6983267 are similar across Africa.

All three branches contain multiple loci with PBS scores that are located in the MHC/HLA region on chromosome 6. For example, rs3817963 has the top PBS score for the Senegalese branch (Fig. 5A). This SNP at 6p21.32 has been associated with lung adenocarcinoma (45). The risk-increasing allele at rs3817963 has an allele frequency of 33.9% in Senegal, 12.9% in Ghana, 10.4% in Nigeria, and 8.4% in South Africa (p-values < 0.0001 for pairwise comparisons between Senegal and other countries, two sample Z-test). Another cancer-associated variant that has large allele frequency differences between African populations is rs2294008, located at 8q24.3. This SNP has the second highest PBS score for the South African branch and it has previously been associated with bladder and gastric cancer (46,47). The risk-increasing allele at rs2294008 has an allele frequency of 28.7% in Senegal, 35.7% in Ghana, 28.8% in Nigeria, and 54.8% in South Africa (p-values < 0.0001 for pairwise comparisons between South Africa and other countries, two sample Z-test).

Focusing on CaP-associated loci, we identify loci with large allele frequency differences between populations, as well as loci that have similar allele frequencies for each African population. For example, rs5919432 is a CaP-associated SNP that is located 71kb from the *androgen receptor* gene at Xq12 (48). This SNP has the highest X-linked PBS score in Fig. 5C. Compared to other study sites, we found that South Africans have elevated frequencies of the risk allele at rs5919432 (Fig. 5D). These allele frequency differences contribute to population-level differences in CaP-risk. We found that the risk-increasing T allele at rs5919432 is more common in MADCaP cases than controls (34.2% vs. 32.1%). Although the association between rs5919432 and CaP was originally discovered in European men, this data reveals that rs5919432 has a similar effect in African populations. The genomic region 8q24.21 contains multiple loci that have been associated with CaP in European men, including rs6983267 (10). Although rs6983267 at 8q24.21 is associated with CaP, it does not have large allele frequency differences between African populations (Fig. 5E). We also found that the risk-increasing G allele at rs6983267 is more common in MADCaP cases than controls (98.2% vs. 97.9%). Note that the protective allele at rs6983267 is rare in Africa but moderately common in Europe (the T allele is found at 50.0% in EUR, 1000 Genomes Project data). This pattern suggests that while rs6983267 contributes to continental-level differences in CaP risk, it has only a minimal effect on population-level differences in CaP risk within sub-Saharan Africa.

### Predicted risks of prostate cancer (CaP) in urban African populations

Using the MADCaP Array we tested whether polygenic risks of CaP vary by population. For each individual, risk scores were calculated by counting risk alleles at 141 CaP-associated loci and weighting by effect size. Higher PRS values indicate that an individual has a higher predicted risk of CaP. Fig. 6 compares PRS distributions for seven African study sites as well as European men from the 1000 Genomes Project, and mean PRS values for each population are indicated by filled rectangles. Overall, we find that predicted risks of CaP are much greater for urban African genomes than European genomes (p-value < 2.2 x 10^-16^, Wilcoxon rank-sum test). This continental-level pattern is consistent with public health data (2). Note that differences in predicted CaP risks between European and African populations exceed differences in predicted risk within Africa. Focusing on MADCaP study sites, there is a substantial amount of overlap in the polygenic risk score distributions of different African populations. Despite this similarity, we observe within-continent heterogeneity for the predicted risk of CaP. The rank order of MADCaP study sites from lowest to highest predicted risk of CaP is: HOGGY, KBTH, SU, WITS, 37 MILITARY, UCH, UATH. Individuals from Dakar, Senegal (gold in Fig. 6) have lower predicted risks of CaP than other African study sites. Conversely, individuals from Abuja, Nigeria (dark green in Fig. 6) have higher predicted risks of CaP than other African study sites. Some of these differences are statistically significant: p-values are ≤ 0.046 for HOGGY vs 37 Military, UATH, UCH, and WITS, and p-values are ≤ 0.031 for UATH vs. HOGGY, KBTH, WITS, and SU (pairwise Wilcoxon rank-sum tests). Rare genetic variants with large effect sizes (e.g. rs183373024 and rs1447295) contribute to the wide tails of each PRS distribution in Fig. 6. Taken together, these results suggest that allele frequency differences at common disease-associated loci can contribute to population-level differences in CaP risk.

**Figure 6.**
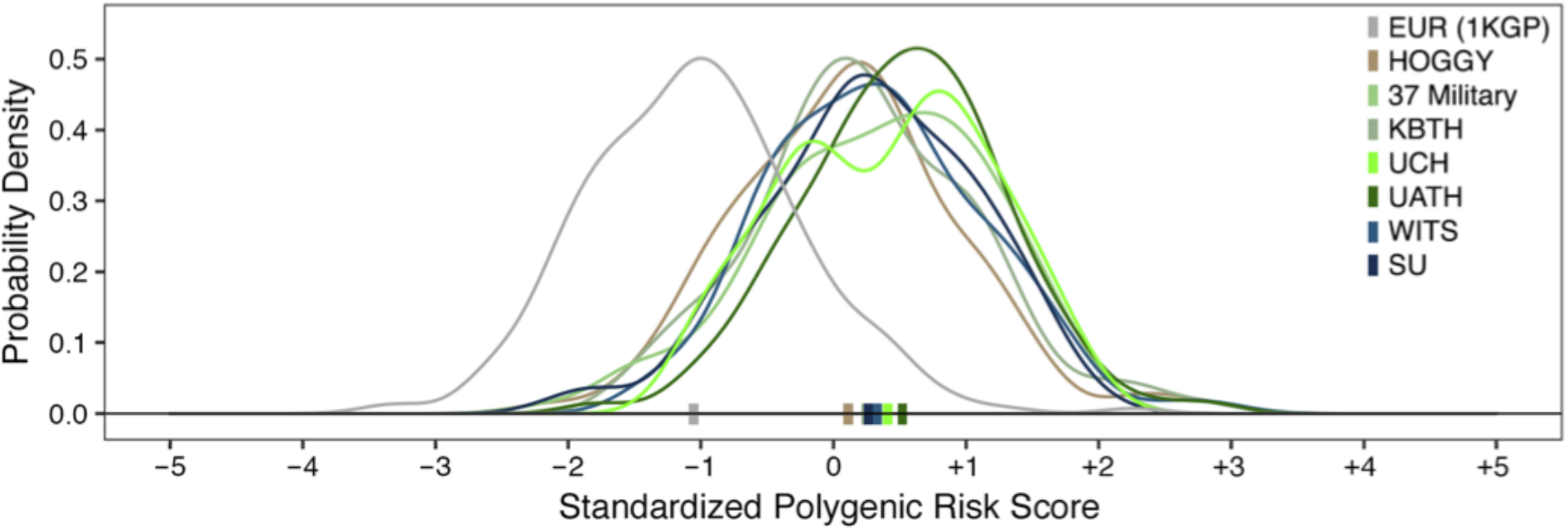
Polygenic risk scores for prostate cancer differ for European and African genomes. Distributions of the genetic risk of CaP are shown for Europeans from the 1000 Genomes Project and Africans from MADCaP study sites. Mean PRS values for each study site are represented by colored rectangles. Markers used in PRS calculations are listed in Table S2.

## Discussion

Using the Axiom genotyping solution, the MADCaP Network has developed a two-peg array that is optimized for studying the genetic basis of CaP in men of African descent. This array successfully tags common and rare variation in African genomes (Fig. 1). The MADCaP Array combines the strengths of the Infinium OncoArray and the H3Africa Array, while maintaining excellent genotyping metrics for diverse African samples. Markers on the MADCaP Array will enable novel disease associations to be discovered and existing cancer associations to be fine-mapped. The 1.5 million markers described in Table S1 are also likely to be of use to researchers developing their own custom genotyping arrays. Applying the MADCaP Array to over 800 African samples, we are able to infer details of population structure, identify loci that contribute to population-level differences in cancer susceptibility, and generate personalized predictions of CaP risk. These findings demonstrate that the MADCaP Array is an effective technology for inferring the population genetics of cancer risks in sub-Saharan Africa.

Sub-Saharan Africa contains substantial amounts of genetic diversity (33,38,49), and this contributes to population-level heterogeneity in cancer risks. For the urban study sites analyzed here, we found that genomes tend to fall into three distinct clusters (Fig. 3C). These clusters broadly match geography: samples from Senegal display similar genetic profiles, samples from Ghana and Nigeria cluster together, and samples from different locations in South Africa cluster together. We also found evidence that the genomes of African individuals contain mixtures of divergent genetic ancestries (Fig. 3E) and that South African study sites have larger effective population sizes than West African study sites (Fig. 4). Clearly, a one-size-fits-all approach is suboptimal when it comes to the genetics of African populations. The genetic heterogeneity of African populations calls for genotyping arrays that accurately capture African polymorphisms.

Genetic risks of cancer have changed during recent human history (50), and our analysis of urban African genomes found many cancer-associated loci that have divergent allele frequencies (Fig. 5, Table S2, and Table S3). There are multiple evolutionary reasons why allele frequencies at cancer-associated loci can differ across human populations. These evolutionary causes include neutral processes like genetic drift and population bottlenecks. Natural selection can also contribute to large allele frequency differences between populations, either directly or indirectly via genetic hitchhiking (20). Regardless of the specific evolutionary cause, differences in allele frequencies at cancer-associated loci can lead to population-level differences in disease risks, as observed in Fig. 6. As SNP-based heritability is a function of allele frequency, loci that are important to disease risks in one population need not contribute much to SNP-based heritability in other populations. Africa is not monomorphic when it comes to the genetic risk of CaP and there is a clear need to conduct studies that cover a broad range of populations.

Genotyping tools such as the MADCaP Array will enable novel cancer associations to be discovered in historically understudied African populations. Smaller LD blocks in African populations will also aid in fine mapping of disease associations. Only by genotyping diverse study cohorts can researchers assess how well polygenic predictions of cancer risks are able to be generalized from large European study cohorts to the rest of the world.

## Author’s Contributions

### Conception and design

T.R. Rebbeck, J. Lachance

### Development of methodology

J. Lachance

### Acquisition of data (provided animals, acquired and managed patients, provided facilities, etc.)

P. Fernandez, M. Jalloh, S.M. Gueye, N.Y. Snyper, B. Adusei, J.E. Mensah, A.O.D. Abrahams, A.O. Adebiyi, A. Orunmuyi, O.I. Aisuodionoe-Shadrach, M.M. Nwegbu, M. Joffe, W.C. Chen, H. Irusen

### Analysis and interpretation of data (e.g., statistical analysis, biostatistics, computational analysis)

M. Harlemon, O. Ajayi, P. Kachambwa, M.S. Kim, C.N. Simonti, M.H. Quiver, A. Mittal, C. Warren, M.H. Woehrmann, P. Zhang, C. Ongaco, E Pugh

### Writing, review, and/or revision of the manuscript

M. Harlemon, O. Ajayi, P. Kachambwa, D.C. Petersen, A.W. Hsing, I. Agalliu, S. Baichoo, A.O. Adebiyi, A. Orunmuyi, O.I,T.R. Rebbeck, L. Petersen, J. Lachance

### Administrative, technical, or material support (i.e., reporting or organizing data, constructing databases)

A.I. Neugut, A.W. Hsing, Y. Quintana, M. Mawhinney, C. Andrews, J. McBride, M. Adams

### Study supervision

L. Petersen, J. Lachance

### Other (performed genotyping)

M. Seutloali, M Fadipe, J. McBride

## Acknowledgments

This work is a product of the MADCaP network (<u>M</u>en of <u>A</u>frican <u>D</u>escent, <u>C</u>arcinoma of the <u>P</u>rostate, https://www.madcapnetwork.org/). This work was supported by a large multi-site NIH/NCI grant (U01CA184374). Additional funding for this work includes startup funds from the School of Biological Sciences at Georgia Institute of Technology to J. Lachance and a seed grant from the Integrated Cancer Research Center at Georgia Institute of Technology.

## Supplemental Data

**Figure S1.**
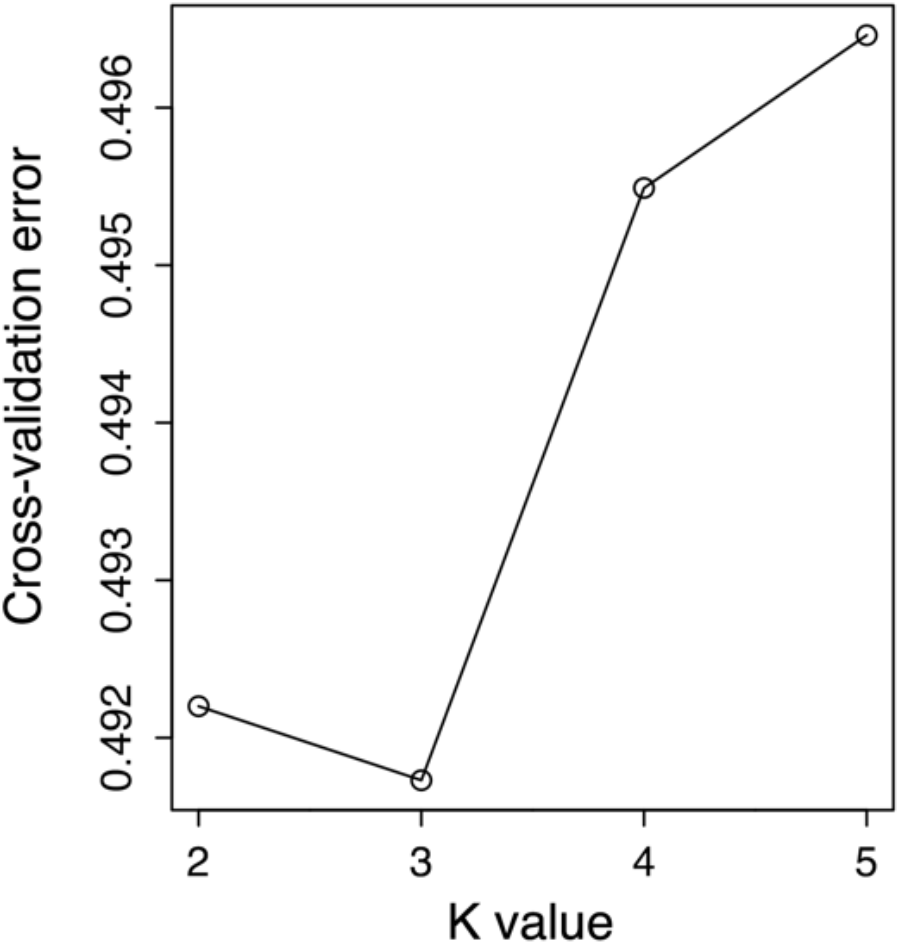
Cross-validation error in ADMIXTURE analyses. The best fit to data occurs at K = 3.

**Table S1. Successfully called markers on the MADCaP Array**. Tab-delimited file that includes genomic positions, inclusion criteria, and overlap with other arrays.

**Table S2. Markers used in PRS calculations**. This list includes all markers from the Schumacher et al. 2018 CaP PRS (10) as well as proxy markers that are found on the MADCaP Array. Chromosome and positions (build hg38) listed here are for MADCaP markers used in PRS calculations. Allele frequencies and PBS scores are also included for the 141 CaP markers used in MADCaP PRS calculations.

**Table S3. Comprehensive list of cancer-associated loci with African allele frequencies and PBS scores**. This dataset includes a total of 2,477 unique markers yielding 5,337 disease or trait associations.

